# Unraveling the effects of spatial variability and relic DNA on the temporal dynamics of soil microbial communities

**DOI:** 10.1101/402438

**Authors:** Paul Carini, Manuel Delgado-Baquerizo, Eve-Lyn S. Hinckley, Hannah Holland-Moritz, Tess E Brewer, Garrett Rue, Caihong Vanderburgh, Diane McKnight, Noah Fierer

## Abstract

Few studies have comprehensively investigated the temporal variability in soil microbial communities despite widespread recognition that the belowground environment is dynamic. In part, this stems from the challenges associated with the high degree of spatial heterogeneity in soil microbial communities and because the presence of relic DNA (DNA from non-living cells) may dampen temporal signals. Here we disentangle the relationships among spatial, temporal, and relic DNA effects on bacterial, archaeal, and fungal communities in soils collected from contrasting hillslopes in Colorado, USA. We intensively sampled plots on each hillslope over six months to discriminate between temporal variability, intra-plot spatial heterogeneity, and relic DNA effects on the soil prokaryotic and fungal communities. We show that the intra-plot spatial variability in microbial community composition was strong and independent of relic DNA effects with these spatial patterns persisting throughout the study. When controlling for intra-plot spatial variability, we identified significant temporal variability in both plots over the six-month study. These microbial communities were more dissimilar over time after relic DNA was removed, suggesting that relic DNA hinders the detection of important temporal dynamics in belowground microbial communities. We identified microbial taxa that exhibited shared temporal responses and show these responses were often predictable from temporal changes in soil conditions. Our findings highlight approaches that can be used to better characterize temporal shifts in soil microbial communities, information that is critical for predicting the environmental preferences of individual soil microbial taxa and identifying linkages between soil microbial community composition and belowground processes.

**Importance:** Nearly all microbial communities are dynamic in time. Understanding how temporal dynamics in microbial community structure affect soil biogeochemistry and fertility are key to being able to predict the responses of the soil microbiome to environmental perturbations. Here we explain the effects of soil spatial structure and relic DNA on the determination of microbial community fluctuations over time. We found that intensive spatial sampling is required to identify temporal effects in microbial communities because of the high degree of spatial heterogeneity in soil and that DNA from non-living microbial cells masks important temporal patterns. We identified groups of microbes that display correlated behavior over time and show that these patterns are predictable from soil characteristics. These results provide insight into the environmental preferences and temporal relationships between individual microbial taxa and highlight the importance of considering relic DNA when trying to detect temporal dynamics in belowground communities.

## Introduction

Information on the temporal dynamics of microbial communities over different time scales can be used to better understand the factors influencing the structure of microbial communities and their contributions to ecosystem processes. The microbial communities found in the human gut (1), leaf litter (2), marine (3), and freshwater (4) habitats can exhibit a high degree of temporal variation. Although the magnitude and timing of this temporal variation in community composition can vary depending on the environment and taxon in question, such temporal variability is often predictable from environmental factors (5). For example, ocean microbial communities display predictable periodic oscillations over time (seasonality) that have been linked to regular changes in biotic and abiotic factors, including phytoplankton dynamics and physicochemical factors (reviewed in refs (3, 6)). These changes in environmental conditions influence the nature of biotic interactions within these ecosystems and can have important ramifications for understanding the functional attributes of microbial communities and the ecosystem services they provide (7–9).

Understanding how temporal changes in environmental conditions influence soil microbial communities is necessary to accurately understand how microbial communities contribute to soil processes and for using microbes as bio-indicators of changes in belowground conditions such as carbon and nutrient availability – parameters that are often difficult to measure directly. However, results from previous studies of temporal variability in soil microbial communities are idiosyncratic. While some studies show soil microbial communities exhibit measurable temporal variation in response to experimental warming (10, 11) and seasonal patterns in temperature and moisture (12–16), other studies show no or minimal variation over time, despite marked changes in environmental conditions (5, 17, 18). One possible explanation for the discrepancies across studies is that the spatial heterogeneity in soil microbial communities—even across short distances—can be sufficiently large to obscure temporal patterns. This hypothesis is supported by numerous studies demonstrating that the spatial variability in soil microbial communities (even across locations only a few meters apart) can be large (for example, ref. (19)). Another explanation is that relic DNA—legacy DNA from dead microbes that can persist in soil—may dampen the observed temporal variability by effectively hiding the true temporal dynamics of soil microbial communities. Relic DNA is abundant in soil (20, 21), and models suggest that during microbial community turnover, relic DNA can mask changes in community structure (21).

We conducted a six month study aimed at disentangling the spatial and relic DNA effects on temporal dynamics in belowground microbial communities. Our study sites were soils on opposing hillslope aspects of a montane ecosystem in the Boulder Creek Critical Zone Observatory (BcCZO) located within the Colorado Front Range of the Rocky Mountains. We intensively sampled two 9 m × 9 m plots, divided into 3 m × 3 m sub-plots every 43-50 days from November 2015 to May 2016 (Fig. 1; five time points total). We chose these locations because the soil microbial communities on the two hillslopes are compositionally distinct (20), relic DNA is abundant (40-60% of the total soil DNA pool, ref. (20)), and the two sites undergo distinct changes in moisture and temperature during this time span (22), providing us with naturally contrasting systems in which to investigate temporal dynamics in belowground microbial communities. We characterized the microbial communities at each site using 16S rRNA gene and internal transcribed spacer 1 (ITS) marker sequencing to profile the prokaryotic (bacterial and archaeal) and fungal communities, respectively. Here, we unravel the relationships between spatial and temporal variability in soil microbial community composition and show the influence of relic DNA on these sources of variability. Further, we use this information on temporal dynamics to identify groups of microbes that share temporal patterns and similar responses to changes in environmental conditions, information that provides insight into the ecologies of understudied soil microbial taxa.

**Figure 1:**
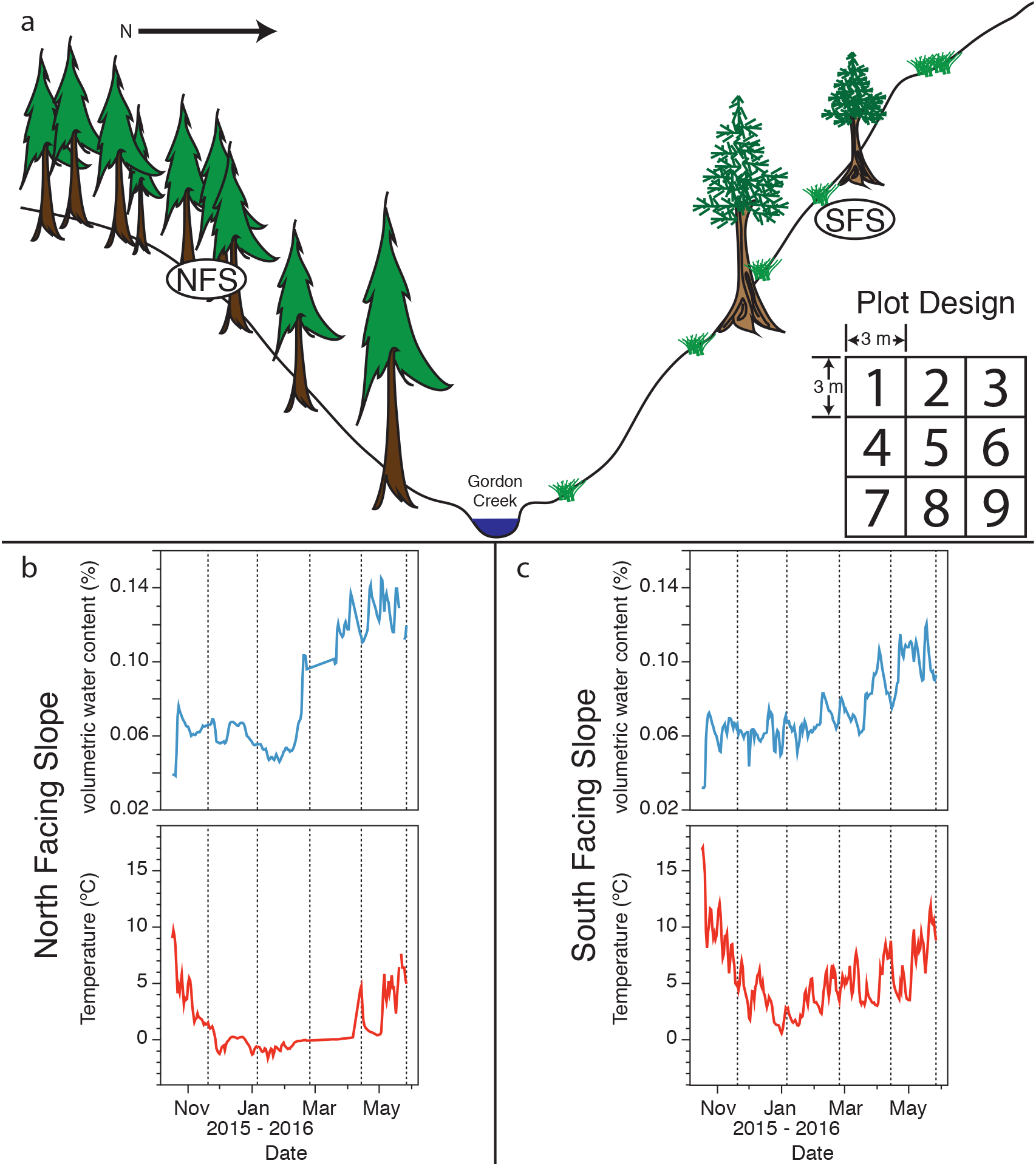
Overview of the Gordon Gulch sampling sites and environmental conditions across the sampling sites. (a) Conceptual diagram of sampling site location and plot design, reproduced with modification from ref. (29). The North facing slope (NFS) plot was centered at 40°0’44.759”N 105°28’9.123”W. The South facing slope (SFS) plot was centered at 40°0’48.551”N 105°28’8.355”W. Inset in (a) is an illustration of plot design. A single plot is comprised of nine 3 m × 3 m replicate sub-plots, as described in the main text. Daily mean soil volumetric water content and soil temperature from *in situ* sensors at 5 cm depth for the NFS (b) and SFS (c) during the course of the experiment. Vertical dashed lines in (b) and (c) indicate sampling dates.

## Results & Discussion

### Spatial variation in soil microbial communities is stronger than temporal variation and is unaffected by relic DNA

Consistent with a previous study conducted at these sites (20), and other studies describing the spatial variability of soil microbial communities (19), the prokaryotic and fungal communities on the south-facing hillslope (SFS) were distinct from those on the north-facing hillslope (NFS), regardless of the time point sampled or whether relic DNA was removed (Supplementary Fig. 1). Most notably, the SFS had higher relative abundances of the archaeal phylum Crenarchaeota (all of which were classified as probable ammonia-oxidizing ‘*Candidatus* Nitrososphaera’), and the bacterial phyla Nitrospirae and Verrucomicrobia (Supplementary Fig. 2). Beyond these expected slope-specific differences, we observed significant intra-plot spatial heterogeneity in microbial community composition that persisted throughout the course of the experiment, and this intra-plot heterogeneity was evident irrespective of whether relic DNA was removed. Before removing relic DNA, there was significant spatial variability across the sub-plots in both prokaryotic and fungal communities on the NFS (Fig. 2 a,e; PERMANOVA R^2^=0.228 and R^2^=0.311; P≤0.001, respectively). These significant spatial differences were still apparent on the NFS for both prokaryotes and fungi after relic DNA was removed (Fig. 2 c,g; PERMANOVA R^2^=0.234 and R^2^=0.292; P≤0.001, respectively). We also found significant spatial variability on the SFS in samples that were not treated to remove relic DNA, but this spatial effect was much more pronounced than on the NFS, with a clear partitioning between sub-plots 5, 6, 8 and 9 (see ‘Plot Design’ in Fig. 1a for numbering) from the remainder of the sub-plots (Fig. 2 b,f; PERMANOVA R^2^=0.308, P≤0.001 for prokaryotes and R^2^=0.317, P≤0.001 for fungi). Similar to the NFS, these strong spatial patterns remained after relic DNA was removed (Fig. 2 d,h; PERMANOVA R^2^=0.310 for prokaryotes and R^2^=0.291 for fungi; P≤0.001). These data show that pronounced spatial variability in soil microbial community composition at the meter scale persists over time. The presence of relic DNA does not affect our overall ability to detect this persistent spatial variation.

**Figure 2:**
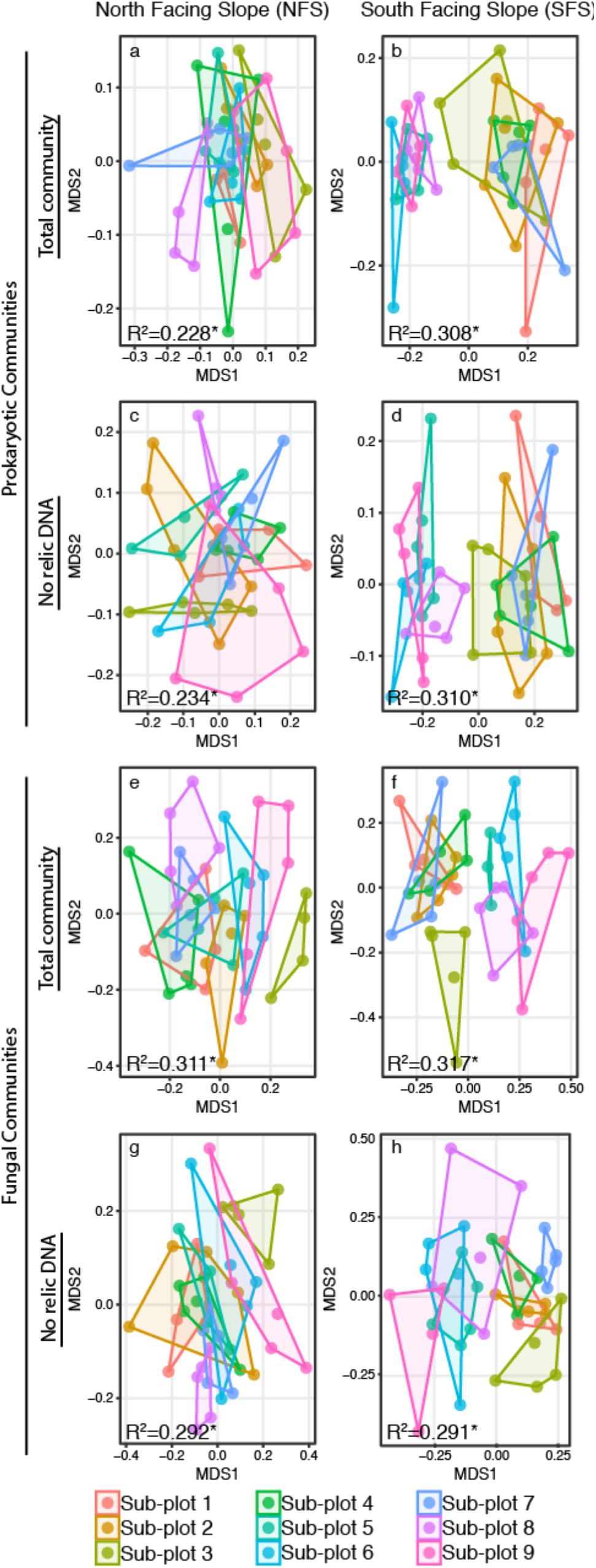
Intra-plot spatial variability in soil microbial communities persists over time on both slopes regardless of whether relic DNA is removed. MDS plots showing the prokaryotic (a-d) or fungal (e-h) communities on the north facing slope (a,c,e,g) and south facing slope (b,d,f,h). Points are colored by sub-plot number (plot layout is illustrated in Fig. 1). Hulls connect the outermost points on each slope. PERMANOVA R^2^ values are listed on each panel; a single asterisk indicates a PERMANOVA P value ≤0.001.

### Removing relic DNA enhanced our ability to detect temporal changes in soil microbial communities

We investigated the effect of relic DNA on temporal variability in belowground microbial communities on a sub-plot basis to control for the aforementioned high degree of intra-plot spatial variability and discriminate between temporal and spatial sources of variation in microbial community structure. When limiting PERMANOVA permutations to within sub-plots over time, we found significant temporal variability for both prokaryotes and fungi on both slopes in untreated control soils (PERMANOVA R^2^ =0.128 P≤0.001 for prokaryotes and R^2^ =0.124 P≤0.001 for fungi on the NFS; and R^2^ =0.110 P≤0.001 for prokaryotes and R^2^ =0.101 P≤0.001 for fungi on the SFS) and in soils that were treated to remove relic DNA (PERMANOVA R^2^ =0.119 P≤0.001 for prokaryotes and R^2^ =0.103 P≤0.001 for fungi on the NFS; and R^2^ =0.098 P≤0.001 for prokaryotes and R^2^ =0.106 P≤0.001 for fungi on the SFS). We found no significant interaction between temporal variability and relic DNA dynamics, suggesting that the presence of differences in microbial community composition between time points is not dependent on the removal of relic DNA. However, on average, the prokaryotic communities on both slopes were significantly more dissimilar over time after relic DNA was removed, compared to untreated control soils that contained relic DNA (Fig. 3; Kruskal-Wallis test *P*≤0.05). These results show that, while compositional differences between time points can be identified in the presence of relic DNA, the removal of ‘legacy’ DNA from dead microbes significantly enhances the ability to detect important temporal variation in the composition of soil prokaryotic communities. While the differences in temporal variability across all sub-plots are significant for the prokaryotic communities in this study, we did observe a similar pattern for fungal communities, albeit not as strong (Fig. 3).

**Figure 3:**
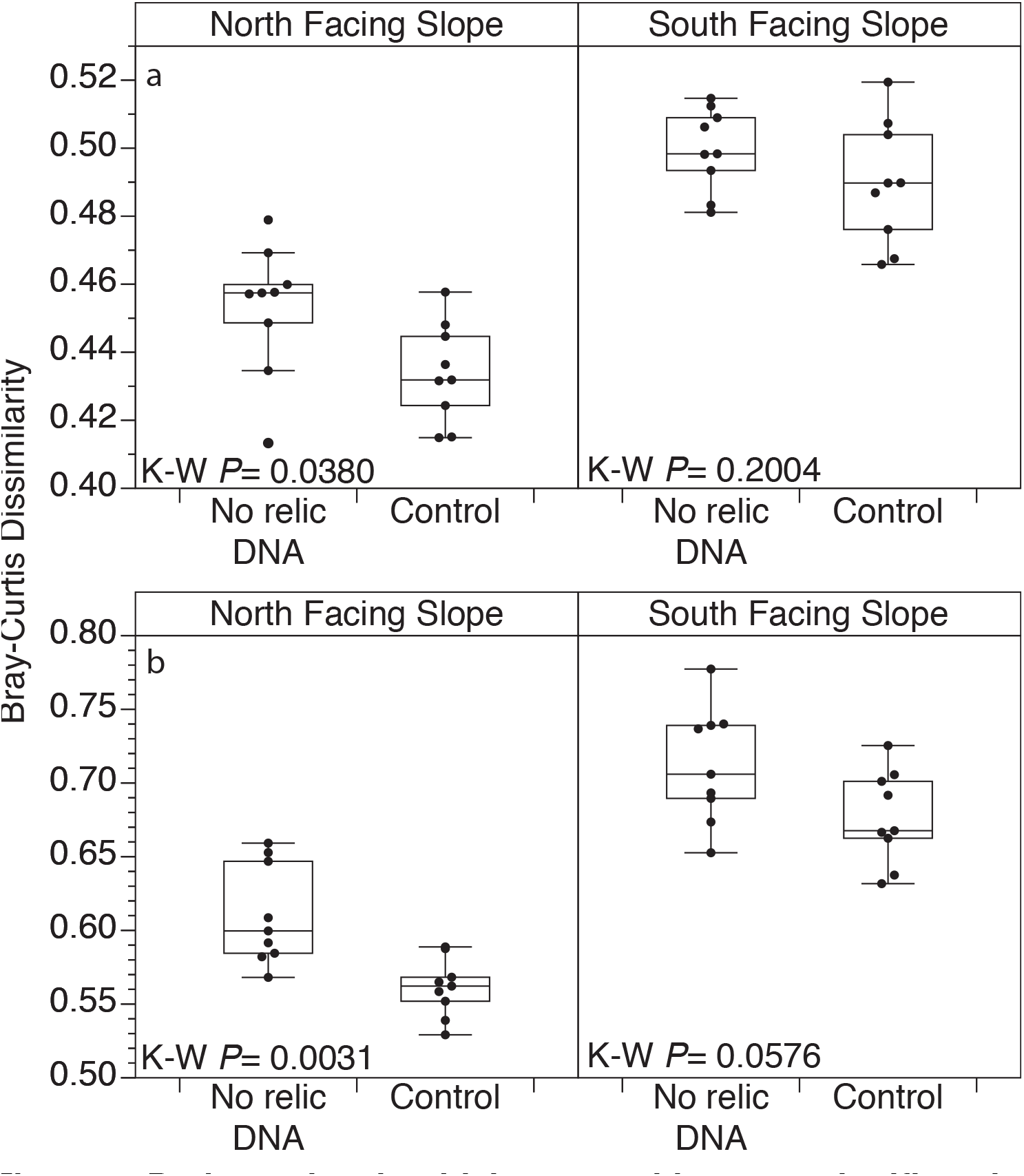
Prokaryotic microbial communities were significantly more dissimilar over time in soils without relic DNA. (a) Prokaryotes (b) Fungi. Points are the mean community dissimilarity for a given sub-plot across all time points (n=5) for samples after relic DNA removal (no relic DNA) or untreated samples (control). Box plots illustrate interquartile range ± 1.5 × interquartile range. The horizontal line in each box plot is the median. Outliers (>1.5 × interquartile range) are shown as points outside of whiskers. Kruskal-Wallis test (K-W) P values are shown.

### Temporal variability in distinct assemblages of prokaryotes and fungi are predictable from soil variables

Characterizing shifts in the relative abundances of individual microbial taxa in temporally dynamic soil systems can give important insight into the ecologies of individual taxa and the environmental factors that influence belowground communities. Thus, we next sought to identify specific groups of taxa that exhibited correlated changes in relative abundances over time in soils after relic DNA was removed. To do this, we used local similarity analysis (LSA) (23) to identify strong (local similarity score ≥0.7) and significant (q-value ≤0.05) positive pairwise microbe-microbe temporal correlations. We constructed and analyzed networks from these correlations and extracted distinct groups (modules) of microbes from NFS and SFS networks using modularity analysis (24) (Fig. 4). On the NFS, the mean normalized relative abundances of 247 microbial taxa (151 bacteria and 96 fungi) were significantly correlated with at least one other taxon over time (Fig. 4a). These correlated taxa clustered into five modules – the mean normalized relative abundances of all five modules changed significantly with time and displayed distinct temporal trajectories (Fig. 4b). On the SFS, 189 taxa (85 bacteria and 104 fungi) were included in the network, and clustered into six modules (Fig. 4c). The relative abundances of five of these six SFS modules changed significantly with time (Fig. 4d).

**Figure 4:**
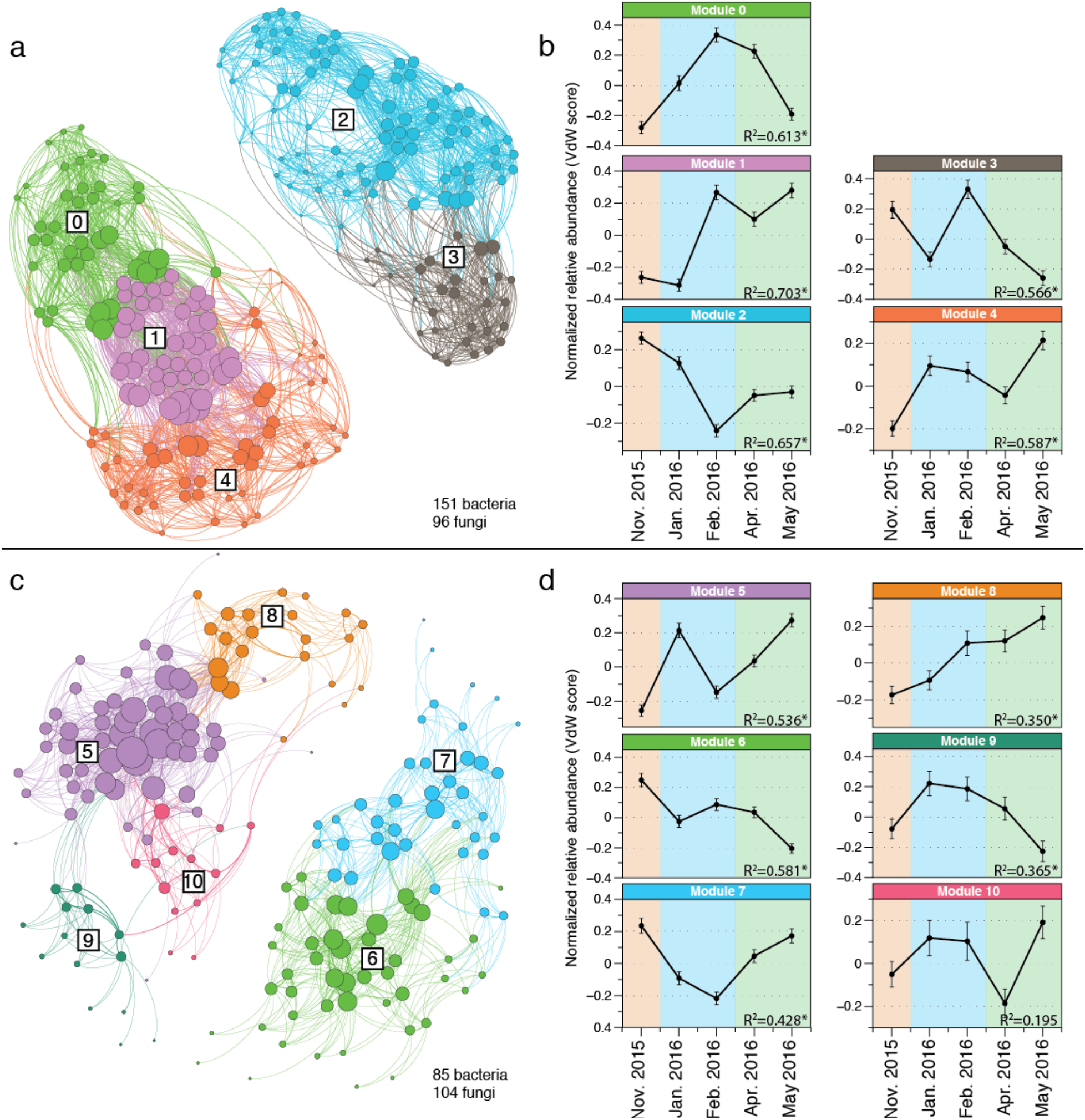
Temporal niche structure in belowground microbial communities. Correlation networks based on significant microbe-microbe temporal correlations for the NFS (a) and SFS (c). Nodes in (a) & (c) are individual prokaryotic or fungal taxa. Lines between nodes represent significant (*q* value ≤ 0.05) and strong (local similarity score ≥ 0.7) positive temporal correlations across all five time points. The sizes of nodes are proportional to the number of correlations to other nodes (the degree), whereby larger nodes have more connections. Colors represent distinct modules, as determined using the modularity algorithm described in ref. (24). Boxed numbers in networks are arbitrary module numbers and match those in panels (b) and (d). Modularity analysis of each network revealed clusters of microbes that have similar temporal patterns. These temporal patterns were plotted for the NFS (b) and SFS (d). Points in (b) and (c) are the mean Van der Waerden (VdW) normalized relative abundance of all taxa in a given module. Error bars show ± SEM. The PERMANOVA P value describing the relationship of the normalized relative abundances in relation to time are shown. P values marked with asterisks are significant at P≤0.005. Background is shaded by season: orange=autumn; blue=winter; green=spring. See Supplementary Table 1 for taxonomic module membership.

A large proportion of the temporal variation in the mean normalized relative abundances of the modules that changed significantly over time could be explained by measured soil or environmental characteristics. At each time point, we measured a suite of soil and environmental parameters, including: snow depth, soil temperature and moisture, extractable inorganic nitrogen (NO_3_- + NH_4_+), salinity (electrical conductivity), extractable phosphorus (P), pH, and the chromophoric properties of water-soluble organic matter (WSOM; a metric of organic matter lability (25)). These measured soil characteristics explained 16 to 56% of the variance in the mean normalized relative abundance of individual modules (Supplementary Fig. 3). Most modules on both slopes were best predicted by climatic variables, most notably soil temperature, soil moisture, and snow depth (Supplementary Fig. 3). These results are in line with previous studies demonstrating how changes in soil temperature (10, 14–16), moisture (26) and snow pack (12) can influence belowground microbial communities. In contrast, module 5 was best explained by changes in inorganic nutrient concentrations (phosphorus; Supplementary Fig. 3). While nitrogen and phosphorus inputs can have predictable (27) and lasting (2) effects on microbial community structure, we have a more limited understanding of how short-term seasonal variation in the availability of these nutrients can influence microbial community dynamics, despite evidence that belowground microbial communities are important mediators of soil nutrient dynamics (28, 29). Our results show that a subset of soil microbes organize into modules that are responsive to these subtle changes in phosphorus availability. Variability in WSOM constituents did not contribute significantly to temporal variability in environmental conditions (Supplementary Fig. 4) and thus, we excluded these measures from the models describing the temporal variability of the modules. Given that previous work at these sites showed a high degree of spatial variation in WSOM distributions (25, 30), we suspect that the pronounced spatial variability in WSOM distributions may have obscured our ability to detect significant effects of WSOM characteristics on the temporal dynamics of the soil microbial communities.

The construction of modules based on shared temporal patterns allowed us to identify biotic or abiotic factors that are correlated with shifts in the relative abundances of individual taxa. As most soil prokaryotic taxa remain undescribed (31), linking the observed temporal dynamics of specific taxa (many of which cannot be classified to the genera or species level of taxonomic classification) to their ecological attributes remains difficult. However, we did identify some bacterial taxa with temporal dynamics that can be explained from our presumed understanding of their ecologies. For example, similar to studies showing *Bradyrhizobium* phylotypes tend to be more abundant in low pH soils (31), we found a single *Bradyrhizobium* phylotype on both slopes (modules 2 and 6) for which pH was a significant predictor (Supplementary Fig. 3), indicating that temporal changes in soil pH influences the relative abundance of this abundant phylotype. Similarly, snow cover was the best predictor for the temporal variability of taxa belonging to module 4 (Fig. 4 and Supplementary Fig. 3). We identified several taxa in module 4 that have been directly linked to the microbial communities associated with snow, including the bacterial phylotypes classified as *Herminiimonas* sp. and *Sphingobacteriaceae* spp. (32, 33) (Supplementary Table 1). We also observed several fungal phylotypes belonging to the Mucorales and Mortierellales orders that clustered in module 4 (Supplementary Table 1). Members of these fungal groups have been termed “snow-molds” and are commonly observed on the surface of soils during snowmelt at these sites (34).

Our study also provides insight into the short-term temporal variation of ectomycorrhizal communities, the environmental factors that influence these patterns and other fungal and prokaryotic taxa that co-vary with ectomycorrhizal fungi. Ectomycorrhizal fungi were found on both slopes and partitioned into several modules that were significantly variable over time (on the NFS, modules 0-4; and on the SFS, modules 6, 7, and 9; Fig. 4 and Supplementary Table 1). On the NFS, 56% of the predicted ectomycorrhizal fungal taxa were found in module 2 (Fig. 4; Supplementary Table 1). Module 2 was best predicted by soil moisture and pH, suggesting that these ectomycorrhizal taxa prefer slightly drier and higher pH soils (Supplementary Fig. 3). However, other ectomycorrhizal taxa on the NFS were best predicted by other combinations of soil characteristics and environmental parameters, suggesting a degree of niche partitioning within soil ectomycorrhizal fungal communities (Supplementary Fig. 3), a finding in agreement with previous observations (35, 36). Fewer ectomycorrhizal taxa displayed correlated behavior with other microbes on the SFS, but the majority of them (69%) belonged to module 6, which was best predicted by snow and moisture. On the SFS, these ectomycorrhizal taxa tended to be more abundant when soils were drier with less snow cover (Supplementary Fig. 3). These findings indicate a degree of temporal niche partitioning in ectomycorrhizal fungal communities on both slopes in response to distinct environmental conditions (Supplementary Fig. 3).

## Conclusions

This study provides new evidence that the temporal dynamics of groups of prokaryotes and fungi living in surface soils are, to some degree, predictable, and that a more detailed characterization of the temporal variability in soil microbial communities is critical to understanding the dynamic nature of the soil microbiome. The extensive spatial and temporal sampling design of our study allowed us to disentangle the relationships between spatial heterogeneity in microbial communities, temporal dynamics of these communities, and the effect of relic DNA on these temporal patterns. Unsurprisingly, spatial variation in community structure at both the hillslope scale, and the meter scale (intra-plot) was the dominant source of variability in this study and relic DNA had no significant effect on our ability to characterize these spatial patterns (Fig. 2 and Supplementary Fig. 1).

When controlling for this spatial variability, we were able to detect significant temporal shifts in microbial community composition, regardless of whether relic DNA was removed or not. We emphasize that the magnitude of the temporal variation in soil microbial communities was consistently lower than the spatial variation, even between sub-plots located only a few meters apart. This spatial variability in surface soil microbial communities was relatively stable over time, suggesting that efforts to describe spatial variation in overall community composition are not necessarily impacted by collecting samples across different time points.

We show that when sites are sampled sufficiently across space, temporal variability is apparent in both soils that have been treated to remove relic DNA and in untreated controls. However, we provide new evidence that the removal of relic DNA results in greater dissimilarity over time, suggesting that by removing relic DNA, we enhance our ability to detect temporal patterns in the belowground communities (Fig. 3). These findings support our previous hypothesis (20), and predictions based on modeling (21), that the presence of relic DNA can dampen temporal patterns in soil microbial communities. The presence of relic DNA, even in high amounts, does not automatically lead to relic DNA biases in other ecosystems (21). However, our data do suggest that relic DNA has important effects on studies of temporal variation in soil microbial communities (and possibly in other ecosystems), and that the consequences of failing to remove relic DNA would not be apparent from single time point samples.

The belowground environment is one of the most complex and dynamic microbial habitats on Earth. By controlling for spatial and relic DNA effects on temporal variability in these soil microbial communities, we identified groups of microbes that have similar temporal dynamics and the environmental factors that predicted their temporal distributions. A deeper understanding of relationships between soil microbiota can help resolve both the roles of individual taxa and potential ‘ecological clusters’ with emergent function. For example, taxa that covary may exhibit similar niche preferences and compete for growth substrates. In contrast, taxa belonging to a given module may broadly respond to similar environmental signals but occupy distinct substrate niches (37). Microbes that are correlated over time may interact through cross-feeding of metabolic substrates or co-utilization of leaky functions (38)—either directly or in a time-lagged manner. Understanding the basis for shared temporal dynamics is important as microbial interactions are crucial in shaping microbial communities (39), but difficult to measure directly (40). Future investigations that combine cell culturing, synthetic microbial communities, and genomics may help resolve the specific drivers of these co-occurrence patterns (37, 41, 42).

## Methods

### Site description, plot design and sampling procedure

The two plots were set up on opposing slopes alongside an instrumented transect at ~2,530 meters elevation (approximately 40.01°N, 105.47°W), chosen on the expectation that there would be a high level of temporal variability in soil microbial communities as a result of intra-annual changes in soil moisture and temperature (22). The north-facing slope (NFS) and south-facing slope (SFS) have distinct soil and vegetation characteristics and experience different water delivery patterns, particularly during snowmelt (22) (Fig. 1). The NFS and SFS soils are Ustic dystrocryept (Catamount series) and Lithic haplstoll, respectively (43). Soil moisture and temperature were variable over the course of the study and followed expected seasonal trends (Fig. 1). In general, the NFS had a higher soil moisture and a lower temperature than the SFS (Fig. 1). The NFS is vegetated with moderately dense *Pinus contorta* (Lodgepole pines) and develops a snowpack during the winter that melts in spring. In contrast, the SFS is much more sparsely vegetated with *Pinus ponderosa* (Ponderosa pines), intervening grasses and *Arctostaphylos uva-ursi* (kinnikinnick) shrubs and experiences pulses of snowmelt throughout the winter and spring. We sampled ~10-15 random soil cores (0-5 cm, mineral soils only; 1” core diameter) within each sub-plot at each of the five time points. The soil cores from each sub-plot were pooled, sieved to 2 mm and homogenized at each time point and partitioned for microbial community and nutrient analyses. Sample dates are reported in Supplementary Table 2.

### Continuous environmental measurements

Several automated measurements were collected every 10 minutes at a meteorological station located near the sample sites (see ‘Data availability’ for data source information). Each slope was instrumented with a soil temperature sensor (Campbell Scientific T-107 temperature probe), and a soil water content reflectometer (Campbell Scientific CS616) located 5 cm below ground. The daily averages from these sensors on each slope are illustrated in Fig. 1b,c. When modelling the relative mean importance of temperature and volumetric water content to module temporal distributions, we used the average of daily mean values from these sensors between sample dates, except for the first time point, which is the mean from the preceding 34 days. Snow depth was measured using digital ultrasonc snow depth sensors (Judd Communications Inc.) fitted with CR1000 dataloggers (Campbell Scientific). Snow depth is reported as mean daily snow depth between sampling points from three sensors on each slope (NFS at snow pole 3, sensors 1-3 and SFS snow pole 10, sensors 9, 11 and 15).

### Discrete environmental measurements

Inorganic N pools were measured for each sub-plot at each time point except for the January 2016 sample on the NFS, sub-plots 1 and 2 and SFS sub-plot 3, where insufficient soil was collected. Sieved soils for inorganic N analyses were stored at 4°C for <72 h. Inorganic N pools were extracted from 10 g field-moist soil in 100 mL 2M potassium chloride with periodic shaking for 18 h and filtered through cellulose Whatman 1 filters. Ammonium (NH_4_+) concentrations were measured in these extracts on a BioTek Synergy 2 with a detection limit of 0.009 mg N L ^−1^ and nitrate (NO_3_-) concentrations were measured on an OI Analytical FS-IV with a detection limit of 0.5603 μg N L^−1^. Dissolved inorganic nitrogen (DIN) was calculated as the sum of NH_4_+ and NO_3_-.

Water-soluble organic matter (WSOM) was analyzed for each sub-plot at each time point except for the following plots, where insufficient sample was collected: NFS February 2016 (all sub-plots); SFS February 2016 sub-plots 1, 8 and 9 and April 2016 sub-plot 5. Sieved soils were stored at −20°C until WSOM extraction. WSOM was extracted by leaching 10 g of soil with 50 ml 0.5 M K_2_SO_4_ following the methods described in (25). The spectroscopically-active portion of the WSOM was characterized with UV-Vis and fluorescence spectroscopy. Samples were diluted to minimize the inner filter effect (44) and the UV-Vis absorbance was measured from 200-800 nm in 1 nm increments using an Agilent 8453 Spectrophotometer with a 1 cm path length. Dissolved organic carbon (DOC) and total nitrogen were measured on a Shimadzu TOC-V. SUVA_254_, a proxy for the aromaticity of the WSOM, was calculated as the absorbance at 254 nm normalized by the DOC concentration (45). Fluorescence scans were collected on a Horiba Jobin Yvon Fluoromax-4 with a 1 cm quartz cuvette and normalized to Raman units (46). The fluorescence index (FI) (47) and humification index (HIX) (48) were calculated from the fluorescence scans using Parallel Factor Analysis (PARAFAC) to further resolve discrete components representing different classes of fluorophores (25).

Other standard soil characteristics were measured at each time point by pooling equal masses of soil from each sub-plot plot on each slope. These measurements included: pH, electrical conductivity (mmhos cm^−1^) and P (ppm). Standard soil chemical analyses were performed at the Colorado State University Soil Water and Plant Testing Laboratory using their standard protocols.

### Relic DNA removal and DNA extraction

Relic DNA was removed as described previously (20). Briefly, 0.03 g of each soil from each sub-plot pool was sub-sampled, resuspended in 3.0 mL phosphate buffered saline (PBS) (1% weight/vol slurry) and either treated with 40 μM propidium monoazide (PMA) in the dark, or left untreated as a control. Both treated and untreated samples were vortexed in the dark for 4 minutes and exposed to a 650-watt light for 4 × 30 s light:30 s dark cycles to activate PMA in treated samples. Light-exposed samples were frozen at −20°C until DNA extraction. DNA was extracted from 800 μL of PMA treated and untreated soil slurries using a PowerSoil-htp 96 well soil DNA Isolation kit (MoBio) following the manufacturer’s instructions, except 770 μL was used in the C2 step. All samples and ‘no soil’ negative controls were randomized into these 96 well DNA extraction plates and extracted simultaneously.

### Amplicon sequencing and analytical methods

For sequence-based analyses of 16S rRNA and ITS marker regions, we used the approaches described previously (20). Briefly, we amplified each sample in duplicate in 25 μl PCR reactions containing: 12.5 μl of Promega GoTaq Hot Start Colorless Master Mix; 0.5 μl of each barcoded primer (10 μM each of bacterial 16S: 515F 5’-GTGCCAGCMGCCGCGGTAA-3’ & 806R 5’- GGACTACHVGGGTWTCTAAT-3’; fungal ITS: 5′-CTTGGTCATTTAGAGGAAGTAA-3′ & ITS2 5′-GCTGCGTTCTTCATCGATGC-3′); 10.5 μl water; 1 μl of template DNA. Thermal cycler program: 94°C for 5 min, followed by 35 cycles of (94°C 45 s; 50°C 60 s; 72°C 90 s) and a final extension 72°C 10 min. Duplicate PCR reactions for each sample were pooled, cleaned and normalized using the ThermoFisher Scientific SequalPrep Normalization Plate kit. Cleaned and normalized amplicons were pooled, spiked with 15% phiX and sequenced on an Illumina MiSeq using v2 500-cycle paired end kits. The samples were sequenced in two batches total – one for prokaryotes and one for fungi. Reads were processed as described in (ref. (27)). Briefly, raw amplicon sequences were demultiplexed according to the raw barcodes and processed with the UPARSE pipeline (49). A database of ≥97% similar sequence clusters was constructed in USEARCH (Version 8) (50) by merging paired end reads, using a “maxee” value of 0.5 when quality filtering sequences, dereplicating identical sequences, removing singleton sequences, clustering sequences after singleton removal, and filtering out cluster representative sequences that were not ≥75% similar to any sequence in Greengenes (for prokaryotes; Version 13_8) (51) or UNITE (for fungi) (52) databases. Demultiplexed sequences were mapped against the *de novo* constructed databases to generate counts of sequences matching clusters (i.e. taxa) for each sample. Taxonomy was assigned to each taxon using the RDP classifier with a threshold of 0.5 (53) and trained on the Greengenes or UNITE databases. To normalize the sequencing depth across samples, samples were rarefied to 10,159 and 7,076 sequences per sample for the 16S rRNA and ITS analyses, respectively. Functional predictions for fungal taxa were obtained using FUNGuild (54).

### Statistical analyses

Calculations of Bray-Curtis dissimilarity by slope, sub-plot and temporal analyses were conducted on the entire 16S rRNA or ITS datasets without filtering. Bray-Curtis distances were calculated on square root transformed taxon relative abundances using the mctoolsr R package (55).

### Temporal analyses and network construction

We identified significant temporal correlations in the relative abundances of individual taxa on each slope that were, on average, ≥ 0.1% of the community across all samples in soils that were treated to remove relic DNA using extended Local Similarity Analysis (eLSA) (23) with the following parameters: lsa_compute -s 5 -r 9 -p perm. We defined significant temporal associations as those with a local similarity (LS) score ≥ 0.7 (i.e.-strong to very strong correlations) and a *q* value ≤ 0.05. Pairs of significantly correlated taxa were analyzed in Gephi (version 0.8.2). Network modularity was calculated by implementing the ‘modularity’ function (24) within Gephi, with a resolution setting of 0.9 for both slopes. Node IDs (individual taxa) belonging to the same module were extracted to delineate temporal patterns in their normalized relative abundances. Normalized relative abundances for each node ID were calculated using the tRank command in the multic R package.

### Random forest analysis

For each slope, we used Random Forest modeling (56) to first identify the measured environmental and soil variables that were significant predictors of time (*P* ≤ 0.05), using time as a response variable (Supplementary Fig. 4). These significant environmental factors are expected to predict changes in module abundance over time. We conducted a second round of Random Forests analysis with the significant environmental predictors shown in Supplemental Fig. 4 to identify the most important environmental factors or soil characteristics that predicted the mean normalized relative abundances of each module (see ref. (57) for a similar approach). The importance (increase in mean square error %) and significance of each predictor was computed for each tree and averaged over the forest (9999 trees) using the rfPermute R package. Significant predictors were defined as those with a *P* value ≤ 0.05. Samples for which environmental and soil characteristics were missing because of insufficient sample were excluded from random forest and Spearman correlation analysis.

## Data Availability

Raw DNA sequence data, the corresponding mapfile and all soil and environmental characteristics are available on figshare.com: 10.6084/m9.figshare.6710087. Snow depth data are available through the Boulder Creek Critical Zone Observatory website: http://criticalzone.org/boulder/data/dataset/2423/. Temperature data for the NFS and SFS are available through the Boulder Creek Critical Zone Observatory website http://criticalzone.org/boulder/data/dataset/2426/.

## Acknowledgements

We thank Gordon Bowman, Youchao Chen, Matt Gebert, Lior Gross, Evan Lih, Emily Morgan, Dillon Ragar, Nathan Rock and Joel Singley for assistance setting up plots, sampling and nutrient analyses. We also thank David Needham for assistance with eLSA and Albert Barberán and Sydney Glassman for critical feedback on previous versions of the manuscript. Funding to support this work was provided by grants from the National Science Foundation EAR 1331828, EAR 1461281, and DEB 1556753 to N.F., and a Visiting Postdoctoral Fellowship award to P.C. from the Cooperative Institute for Research in Environmental Sciences at the University of Colorado.

## Notes

#### Summary of Updates

The manuscript has been revised to analyze five time points instead of time. After the initial submission was prepared, we noticed a DNA extraction batch effect that precluded the analysis of the full dataset. The main findings of the preprint remain unchanged.

