## Supplemental File 1 for "Unraveling the effects of spatial variability and relic DNA on the temporal dynamics of soil microbial communities"

Supplementary information
For

### Unraveling the effects of spatial variability and relic DNA on the temporal dynamics of soil microbial communities

Paul Carini[1,2], Manuel Delgado-Baquerizo[1,3], Eve-Lyn S. Hinckley[4,5], Hannah Holland-Moritz[1,7], Tess E Brewer[1,6], Garrett Rue[4], Caihong Vanderburgh[1], Diane McKnight[4] and Noah Fierer[1,7]

**Supplementary Table 1:** Module membership by slope, including FUNGuild functional predictions for fungi. (see accompanying excel file)

**Supplementary Table 2:** Time points, sampling dates, days between timepoints, and season

| Time point | Date | Days since previous | Astronomical Season |
|---|---|---|---|
| T01 | November 20, 2015 | 0 | Autumn |
| T02 | January 06, 2016 | 47 | Winter |
| T03 | February 25, 2016 | 50 | Winter |
| T04 | April 14, 2016 | 48 | Spring |
| T05 | May 27, 2016 | 43 | Spring |

**Supplementary Figures:**

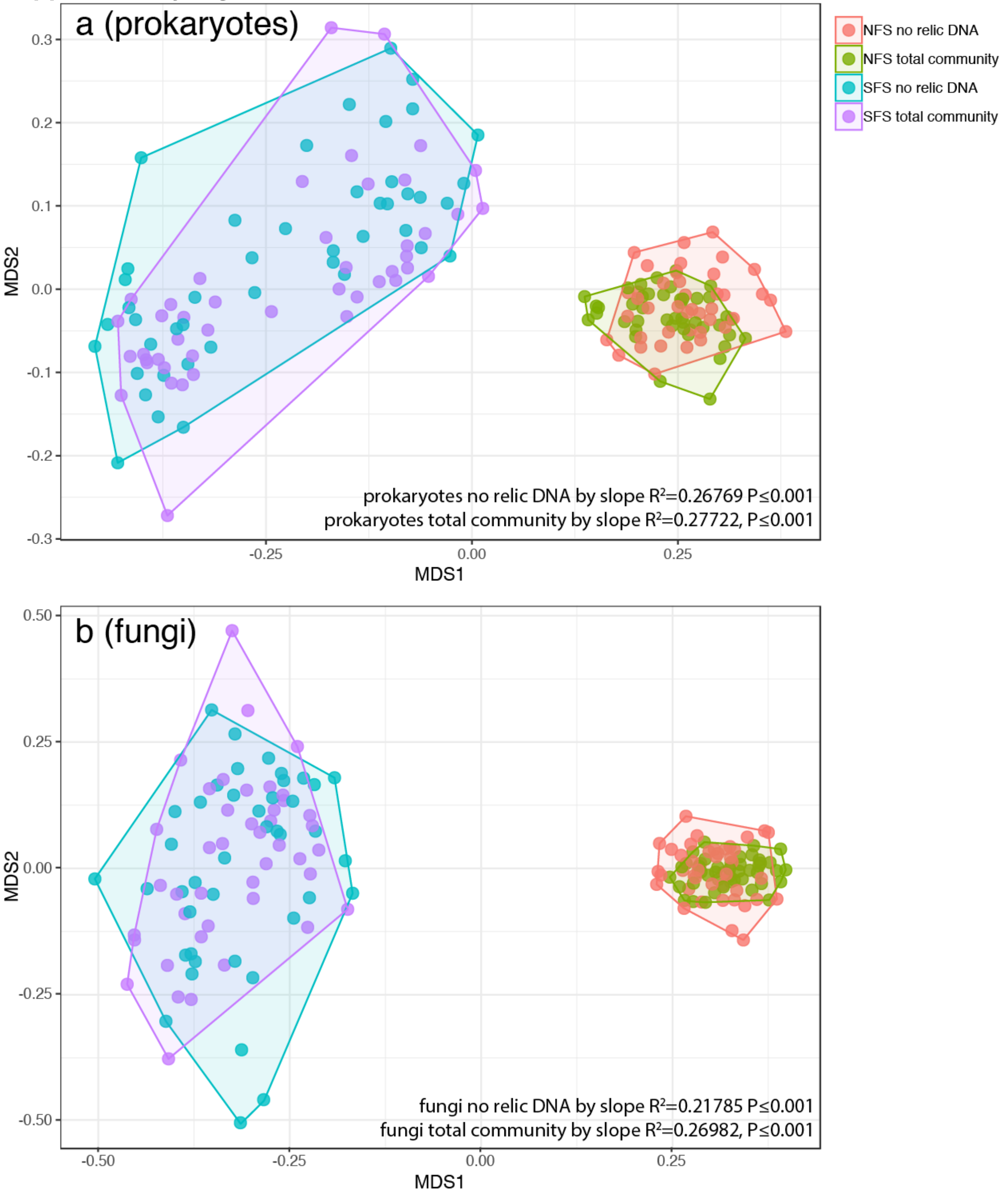


**Supplementary Figure 1: The microbial communities on the NFS are distinct from those on the SFS, regardless of time point sampled or whether relic DNA was removed.** Non-metric multidimensional scaling (NMDS) plot showing the prokaryotic (a) or fungal (b) communities for each sub-plot for each time point on both slopes. Points are colored by slope and whether relic DNA was removed. Hulls connect the outermost points on each slope. Permutational multivariate analysis of variance (PERMANOVA) statistics for slope differences are shown with and without relic DNA removal.

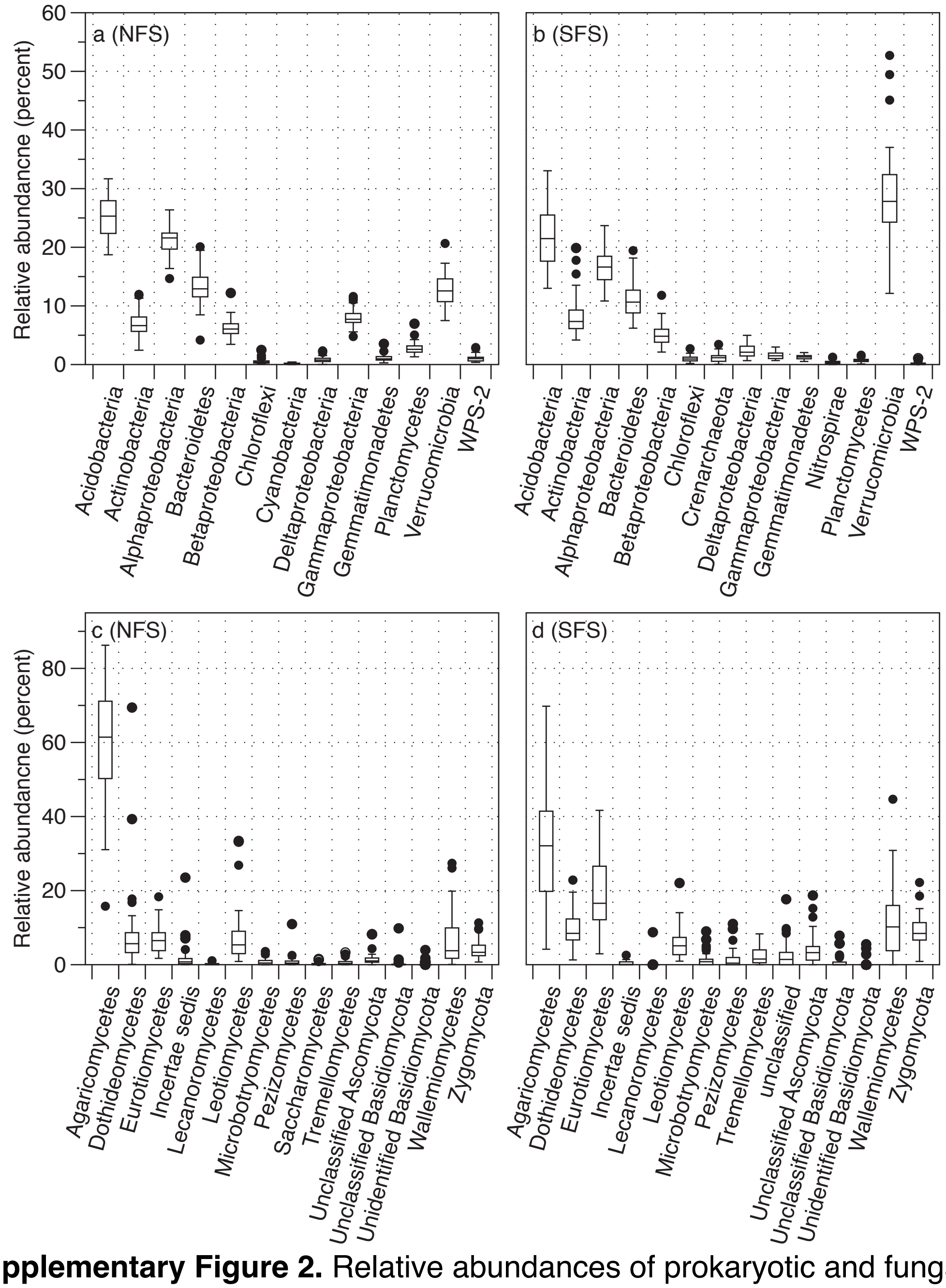


**Supplementary Figure 2.** Relative abundances of prokaryotic and fungal taxa on the north facing slope (NFS) and south facing slope (SFS). Box plots show the distributions of the relative abundances of prokaryotic (a & b) or fungal (c & d) phyla or classes in each sub-plot for each time point. Box plots illustrate interquartile range ± 1.5 × interquartile range. The horizontal line in each box plot is the median. Outliers (>1.5 × interquartile range) are shown as points.

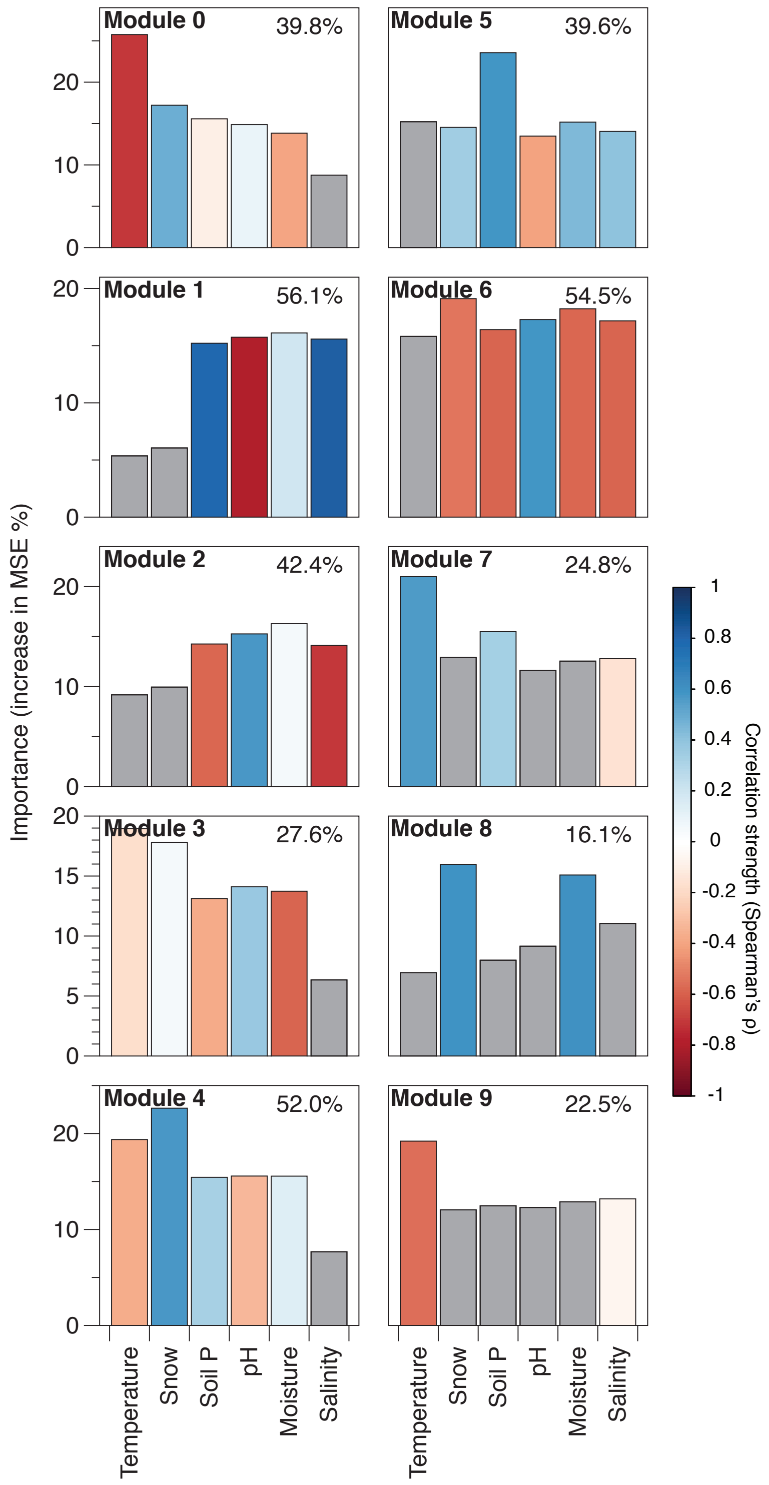


**Supplementary Figure 3: A large amount of the variation in the relative abundances of taxa within modules can be explained by temporal variation in measured soil characteristics.** The percent of temporal variation in the mean normalized relative abundances for each module that is explained by measured soil characteristics is listed in the upper right corner of each plot. Bar heights are the importance (percent increase in mean squared error (MSE)) of measured environmental and soil characteristics to the observed temporal changes in the mean normalized relative abundance of each module as determined by Random Forest modeling. Only environmental variables that changed significantly over time within each slope were included in the models (see Supplementary Fig. 4). Module 10 was excluded from modeling because the normalized relative abundances did not change significantly with time. The color scale indicates the strength and direction of Spearman correlation between the measured environmental variables and the mean normalized relative abundances of the modules. Variables that were not significant predictors (P>0.01) to the Random Forest models are colored grey.

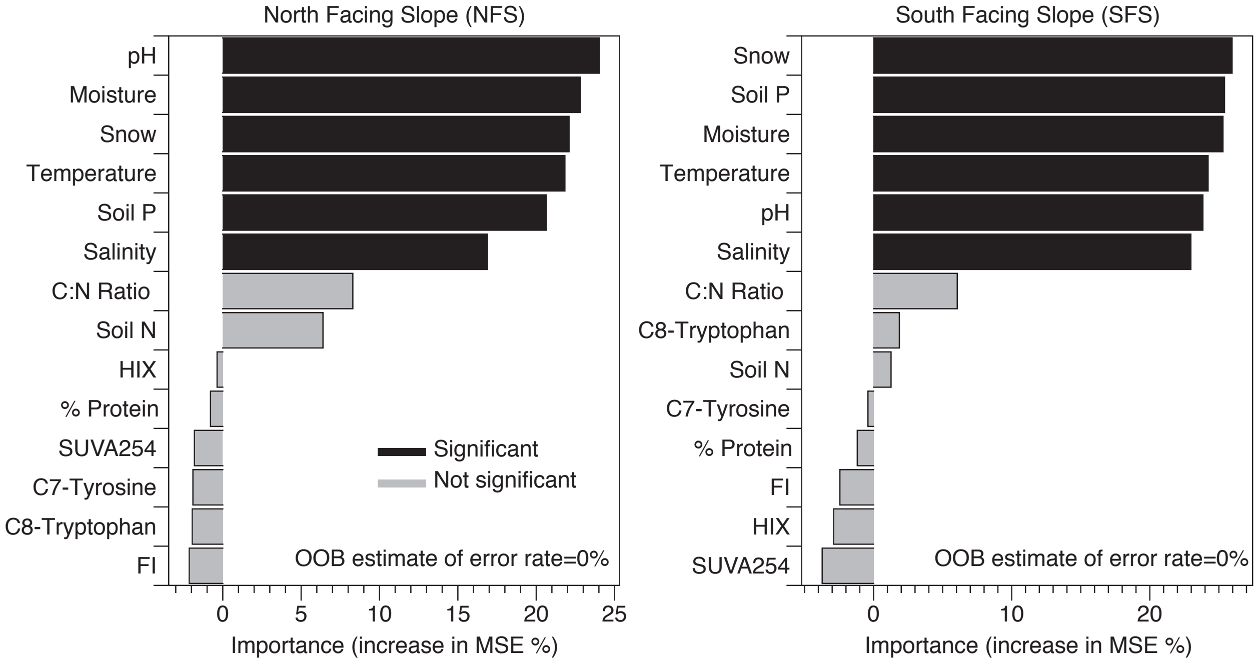


**Supplementary Fig. 4: Results from the random forest analyses that were used to identify key environmental factors significantly correlated with time within each slope.** Time was treated as a categorical variable in analyses.
